# Aβ oligomers induce sex-selective differences in mGluR5 pharmacology and pathophysiological signaling in Alzheimer mice

**DOI:** 10.1101/803262

**Authors:** Khaled S. Abd-Elrahman, Awatif Albaker, Jessica M. de Souza, Fabiola M. Ribeiro, Michael G. Schlossmacher, Mario Tiberi, Alison Hamilton, Stephen S. G. Ferguson

## Abstract

Sex is a key modifier of the prevalence and progression of Alzheimer’s disease (AD). β- Amyloid (Aβ) deposition is a pathological hallmark of AD and aberrant activation of metabotropic glutamate receptor 5 (mGluR5) by Aβ has been linked to AD progression. We find that mGluR5 exhibits distinct sex-dependent pharmacological profiles. Specifically, endogenous mGluR5 from male mouse cortex and hippocampus binds with high-affinity to Aβ oligomers whereas, female mGluR5 exhibits no affinity to Aβ oligomers. The binding affinity of mGluR5 to Aβ oligomer is dependent on its interaction with cellular prion protein (PrP^C^) as mGluR5 co-immunoprecipitates with PrP^C^ from male, but not female, mouse brain. Aβ oligomers also bind with high-affinity to human mGluR5 in male, but not female, cortex. The mGluR5/Aβ oligomer/PrP^C^ ternary complex is essential to elicit mGluR5-dependent pathological signaling and as a consequence mGluR5-regulated GSK3β/ZBTB16 autophagic signaling is dysregulated in male, but not female, primary neuronal cultures. These sex-specific differences in mGluR5 signaling translate into in vivo differences in mGluR5-dependent pathological signaling between male and female AD mice. We show that the chronic inhibition of mGluR5 using a mGluR5-selective negative allosteric modulator reactivates GSK3β/ZBTB16-regulated autophagy, mitigates Aβ pathology and reverses cognitive decline in male, but not female, APPswe/PS1ΔE9 mice. Thus, it is evident that, unlike male brain, mGluR5 does not contribute to Aβ pathology in female AD mice. This study highlights the complexity of mGluR5 pharmacology and Aβ oligomer-activated pathological signaling and emphasizes the need for clinical trials redesign and analysis of sex-tailored treatment for AD.

## INTRODUCTION

Alzheimer’s disease (AD) is a progressive neurodegenerative disease characterized by age-related memory loss and cognitive decline and despite the alarming prevalence, no available treatments exist to either modify or reverse its progression *(1–3)*. β-Amyloid (Aβ) oligomers and hyper-phosphorylated tau protein represent pathological hallmarks of AD and accumulate with disease progression to disrupt neuronal signaling and trigger neurodegeneration *(4–6)*. Sex is an important modulator of AD prevalence and both clinical and preclinical evidence indicate that the incidence in females is higher than age- and risk factor-matched males *(7–10)*. This higher incidence of cognitive impairment is commonly attributed to reduced estrogen and/or estrogen receptors levels following menopause *(11–13)*. However, this assumption was weakened by the first large-scale clinical study demonstrating increased risk of dementia and poor cognitive outcomes following hormone-replacement therapy (Women’s Health Initiative Memory Study, WHIMS) *(14, 15)*.

Glutamate, the main excitatory brain neurotransmitter, plays a key role in learning and memory and the G_αq_-coupled metabotropic glutamate receptors 5 (mGluR5) is of particular interest in AD pathology *(5, 16)*. mGluR5 functions as an extracellular scaffold for Aβ and cellular prion protein (PrP^C^) and the complex between PrP^C^ and mGluR5 is essential for Aβ binding *(17–19)*. Aβ also promotes mGluR5 clustering, increases intracellular Ca^2+^ release and inhibits autophagy *(17, 19, 20)*. Moreover, the pharmacological and genetic silencing of mGluR5 reverses cognitive deficits and reduces Aβ pathology in male AD rodent models *(20–24)*. While these studies emphasized the key role of mGluR5 in AD pathophysiology, it is not yet clear whether alterations in mGluR5 signaling are conserved between male and female AD models. Given that both clinical and experimental evidence indicate a divergent AD pathology between males and female, the dramatic drug treatment failure rate in AD clinical trials warrant a better delineation of the molecular signaling mechanism(s) underlying sex-related pathophysiological differences in AD *(7, 10)*. It is important to note that some previous studies have attempted to pharmacologically target mGluR5 in AD mice from both sexes but surprisingly the results were not stratified according to sex and therefore, an accurate representation for the contribution of pathological mGluR5 signaling to AD in females cannot be reached *(25, 26)*.

Our rationale was to assess whether mGluR5 differentially contributes to pathology in both sexes and therefore, we evaluated the binding affinity of Aβ to mGluR5 and the efficacy of Aβ in triggering pathological autophagic signaling in male and female AD mice. We also tested whether the mGluR5 orally bioavailable negative allosteric modulator (NAM), 2-chloro-4-((2,5-dimethyl-1- (4-(trifluoromethoxy)phenyl)-1H-imidazol-4-yl)ethynyl) pyridine (CTEP) *(27)*, differentially alters cognition and progression of Aβ pathology between male and female APPswe/PS1ΔE9 (APP) mice *(28)*. It is worth noting that although other mGluR5 ligands have been reported to only either improve memory deficits (BMS-984923) or Aβ pathological signaling (CDPPB), but not both, in AD mouse models *(25, 26)*. Our choice of CTEP was based on its superior capabilities in reversing Aβ pathology and improving cognitive function with no evidence of drug-related adverse outcomes after extended treatment regimens *(22, 24)*.

Our findings indicate that Aβ oligomers binds with high affinity to mGluR5 in both male mouse and human cortex, but exhibits no affinity for mGluR5 in either female mouse or human brain tissue. The binding of Aβ oligomers to mGluR5 was PrP^C^-dependent as mGluR5 co-immunoprecipitated with PrP^C^ from male, but not female, mouse hippocampus. When wild-type male E18 embryonic neuronal cultures were treated with Aβ oligomers, it inhibited one of the key mGluR5-regulated autophagic pathways, the GSK3β/ZBTB16 pathway. Specifically, Aβ increased pS9-GSK3β phosphorylation and blocked ZBTB16-mediated autophagy in an mGluR5-dependent manner. In contrast, the treatment of female cultures with Aβ did not alter pS9-GSK3β phosphorylation and had no effect on autophagy. Our in vitro findings were translatable in vivo since treatment of male APP mice with mGluR5 NAM was associated with enhanced ZBTB16-regulated autophagy, reduced Aβ pathology and improved cognitive function. However, CTEP exhibited no efficacy in female APP mice and rather caused cognitive impairment in wild-type female mice. These studies revealed that mGluR5 exhibits unexpected and distinct pharmacological profiles with respect to Aβ oligomer/PrP^C^ interactions in male and female brain that can dictate the relative contribution of mGluR5 to AD pathology and therapeutics between both sexes.

## RESULTS

### Sex-specific interaction of Aβ oligomers and PrP^C^ with mGluR5 in mouse brain

Aβ oligomers binds with high affinity to the PrP^C^/mGluR5 complex and the Aβ oligomer/PrP^C^ complex can then drive the pathological signaling of mGluR5 *(17, 29)*. Therefore, we first tested whether the binding of Aβ oligomers to endogenous mGluR5 is sex-specific. We assessed Aβ oligomer-mediated displacement of radiolabeled mGluR5 antagonist (MPEP) binding to endogenously expressed mGluR5 in plasma membrane preparations from male and female wild-type mouse cortex and hippocampus. To our surprise, Aβ oligomers effectively displaced [^3^H]- MPEP binding to male, but not female mouse cortical (Fig. 1A and 1B and Table1) and hippocampal membrane preparations (Fig. 1C and 1D and Table 1). The displacement of [^3^H]- MPEP binding by non-radioactive MPEP was not different when both sexes were compared in each brain regions (Fig. 1A-D and Table 1). Thus, Aβ oligomers binding assays revealed for the first-time distinct sex-dependent pharmacological profiles for mGluR5.

**Figure 1:**
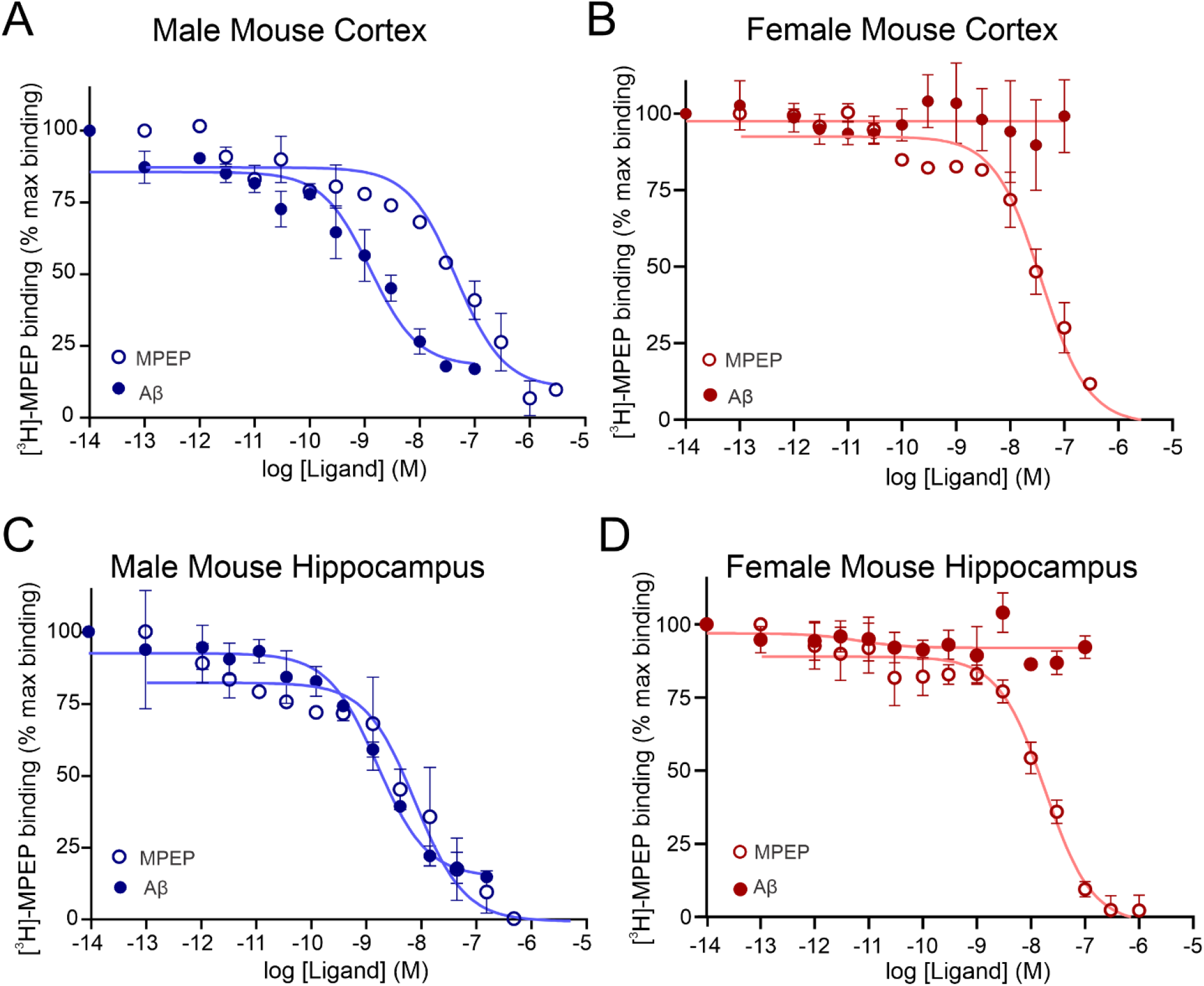
Sex-specific binding of Aβ oligomers to mGluR5 in mouse brain. [^3^H]-MPEP displacement curves for MPEP and Aβ oligomers in membrane preparations from wild type **(A)** male mouse cortex, **(B)** female mouse cortex, **(C)** male mouse hippocampus and **(D)** female mouse hippocampus. Each binding curve represents the mean ± SEM of 3-4 independent duplicate experiments.

**Table 1:** Maximum specific binding and calculated K_i_ for MPEP and Aβ oligomers in mouse and human male and female membrane preparations.

|  |  | Male |  | Female |  |
| --- | --- | --- | --- | --- | --- |
| | | Max. binding (DPM) | $K_i$ (M) | Max. binding (DPM) | $K_i$ (M) |
| Mouse Cortex | MPEP | 6953 ± 156.3 | 4.18E-08 ± 1.48E-08 | 6837 ± 129.3 | 2.27E-08 ± 7.15E-09 |
| | Aβ | 6257 ± 479.1 | 8.46E-11 ± 4.49E-11 | 9601 ± 218.5 | ND |
| Mouse Hippocampus | MPEP | 6872 ± 313.1 | 7.91E-09 ± 6.86E-09 | 6345 ± 166.5 | 9.18E-09 ± 5.74E-10 |
| | Aβ | 7640 ± 307.8 | 6.02E-11 ± 5.05E-12 | 8872 ± 352.4 | ND |
| Human Cortex | MPEP | 5709 ± 116.3 | 9.60E-09 ± 6.57E-09 | 8003 ± 259.8 | 5.04E-09 ± 2.26E-09 |
| | Aβ | 6090 ± 102.5 | 9.78E-11 ± 5.75E-11 | 8025 ± 203.6 | ND |

$K_i$ : Equilibrium dissociation constant; DPM: Disintegrations per minute

Since Aβ oligomers binding to mGluR5 is known to be dependent on the interaction of the receptor with PrP^C^ *(17, 29–31)*, we tested whether PrP^C^ interacts with mGluR5 in a sex-specific manner. We first confirmed that the size of the Aβ oligomers that we employed in our study is capable of interacting with mGluR5/PrP^C^ scaffold and that PrP^C^ is key for Aβ binding to mGluR5 by performing radioligand binding assays in PrP^C^ null (CF10) cells transfected with mGluR5. In control experiments, we found that Aβ oligomers did not displace [^3^H]-MPEP binding in CF10 cells (Fig 2A), but that co-transfection of CF10 cells with PrP^C^ restored the ability of Aβ oligomer to bind mGluR5 (Fig. 2B). We then tested whether PrP^C^ binding to mGluR5 was different between both sexes and found that mGluR5 co-immunoprecipitated with PrP^C^ from male, but not female, wild-type mouse hippocampal tissue (Fig. 2C). Binding specificity of PrP^C^ antibody to mGluR5 was validated by the lack of receptor immunoprecipitation in hippocampus from mGluR5^−/−^ mice. Overall, our findings indicated that, unlike what was observed for males, the mGluR5/ Aβ oligomer/PrP^C^ ternary complex is not formed in female mouse brain.

**Figure 2:**
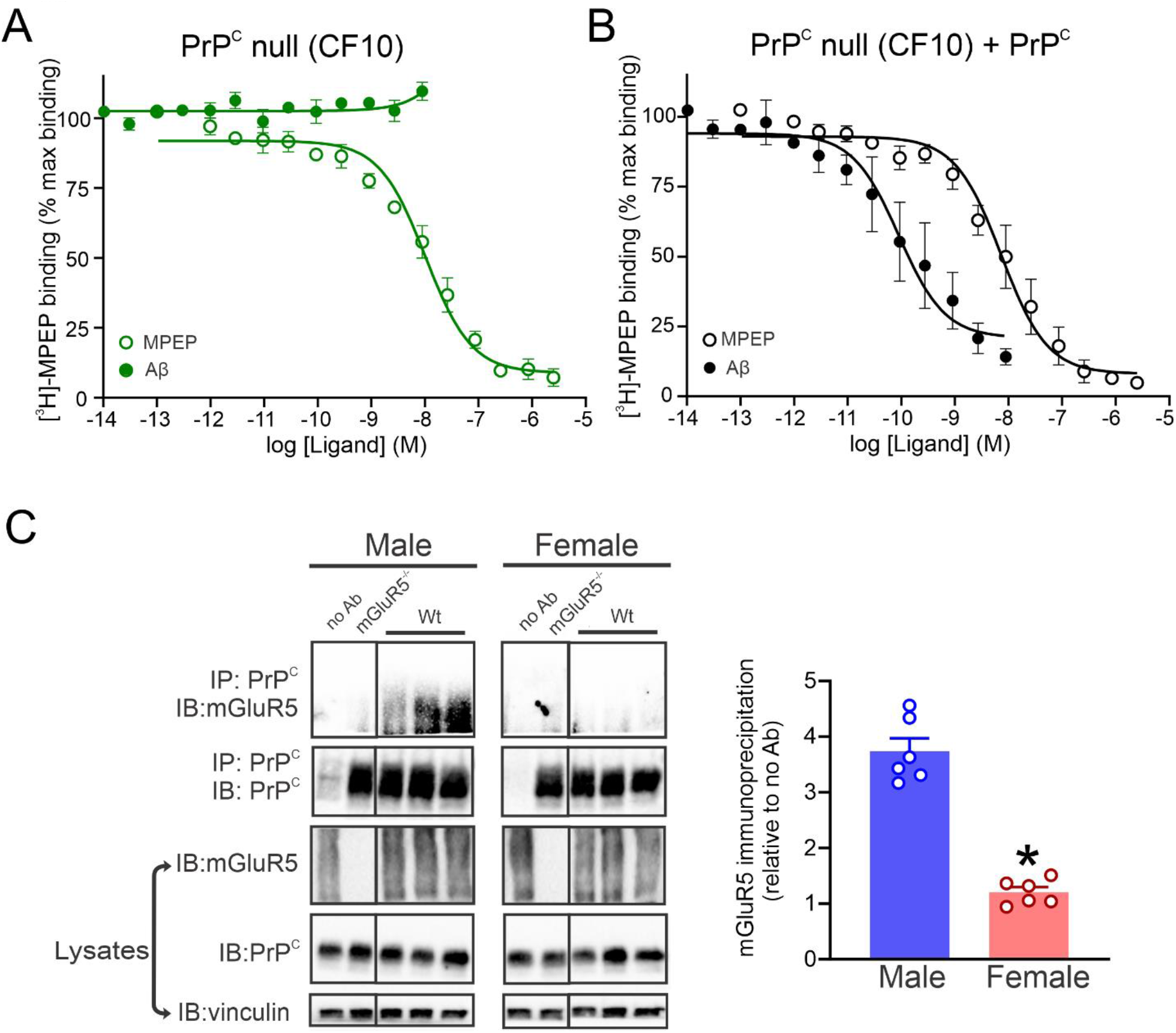
Sex-specific interaction of PrP^C^ with mGluR5 in mouse hippocampus. [^3^H]-MPEP displacement curves for MPEP and Aβ oligomers in membrane preparations from CF10 cells transfected with **(A)** FLAG-mGluR5 or with **(B)** FLAG-mGluR5 + PrP^C^. Each binding curve represents the mean ± SEM of 3 independent duplicate experiments. **(C)** Representative immunoblot (IB) and quantification of mGluR5 co-immunoprecipitated with PrP^C^ (IP) from male and female wild-type (Wt) hippocampus with corresponding lysates. Hippocampal tissue from male and female mGluR^−/−^ or Wt mice in the absence of PrP^C^ antibody (no Ab) served as a negative control (n=6). Immunoprecipitation was calculated relative no Ab lane of each sex.

### Sex-specific binding of Aβ oligomers to mGluR5 in human cortex

To validate that sex-specific binding of Aβ to mGluR5 was not exclusive to mouse tissues, we performed [^3^H]-MPEP binding displacement assays in cortical membranes prepared from male and female human cortical brain samples obtained at autopsy. Identical to what was observed for mouse cortical membranes, displacement of [^3^H]-MPEP binding by MPEP was not significantly different between male and female human cortical membranes (Fig. 3A and 3B and Table 1). However, Aβ oligomers only displaced [^3^H]-MPEP binding to male membranes (Fig. 3A and 3B and Table 1) suggesting that the differential binding of Aβ oligomers to mGluR5 between males and females is an evolutionally-conserved phenomenon and may indeed contribute to gender-specific pathophysiology in human AD patients.

**Figure 3:**
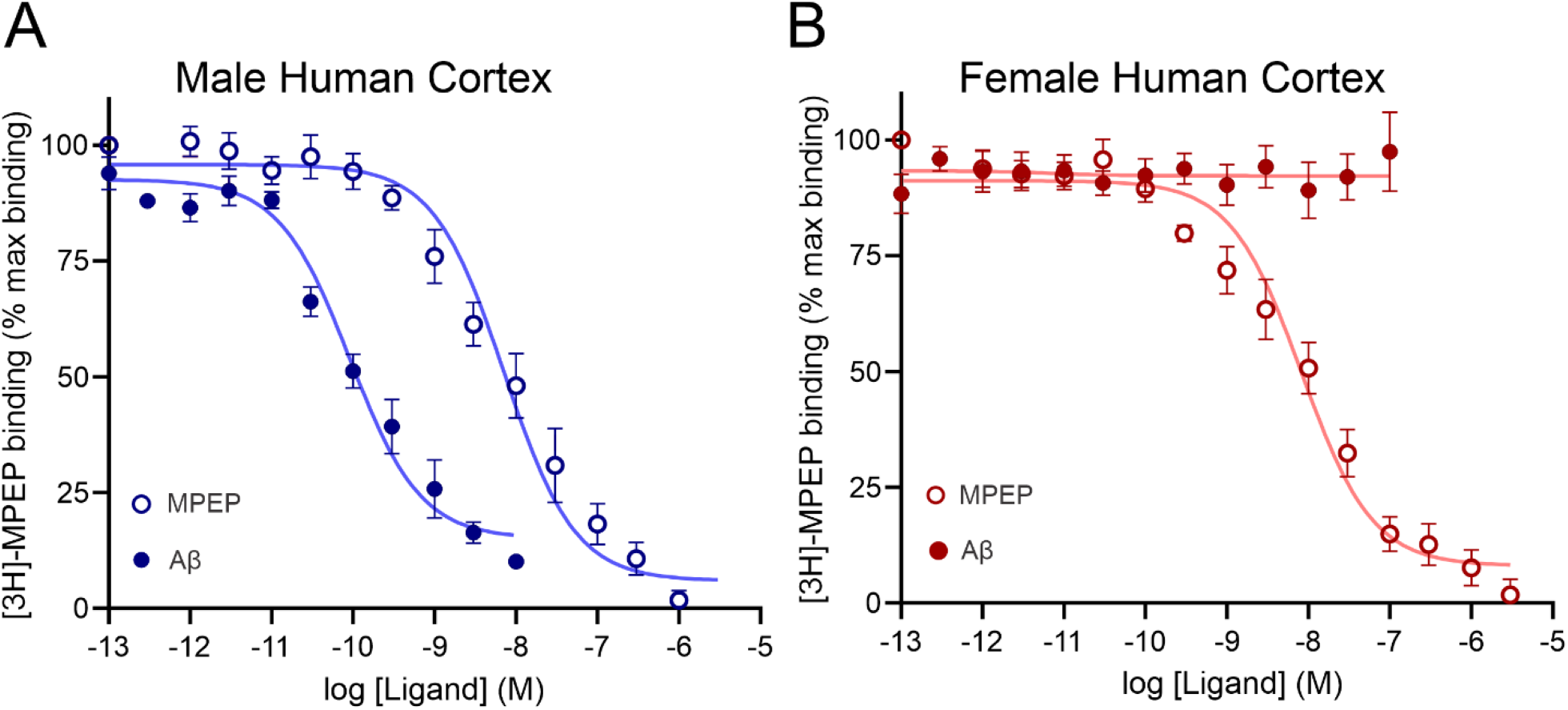
Sex-specific binding of Aβ oligomers to mGluR5 in human brain. [^3^H]-MPEP displacement curves for MPEP and Aβ oligomers in membrane preparations from **(A)** male human cortex, **(B)** female human cortex. Each binding curve represents the mean ± SEM of 4-5 independent duplicate experiments.

### Sex-specific Aβ-activated mGluR5 signaling in primary embryonic neuronal cultures

Since Aβ oligomers binding to mGluR5 was shown to induce receptor clustering and activate its pathological signaling *(17, 19)*, we tested whether Aβ oligomer binding to mGluR5 could inactivate GSK3β/ZBTB16 autophagy pathway, which we identified to be engaged downstream of mGluR5, in a sex-specific manner in the absence of sex hormones in the media. The GSK3β/ZBTB16 pathway was one of the mGluR5 autophagic signaling mechanisms that was previously observed to be dysregulated in male AD mouse models and correlated with a loss of clearance of Aβ *(20, 24, 32)*. Thus, to avoid potentially confounding influence of sex hormones on mGluR5 signaling, we examined Aβ-evoked signaling in primary wild-type male and female E18 mouse cortical neurons in cell culture. Sex of the cultures was determined by PCR amplification to detect X chromosome-linked Rbm31 gene relative to its divergent Y chromosome gametolog *(33)* (Fig. 4A). Treatment of male cultures with Aβ oligomers (100 nM) induced pS9-GSK3β phosphorylation, increased ZBTB16 expression and inhibited autophagy as measured by elevated p62 expression in a manner that was inhibited by CTEP (10 μM) (Fig. 4B). These changes were not observed in cultures derived from female embryos (Fig. 4B). These results clearly demonstrated that Aβ oligomers inactivated ZBTB16 autophagic signaling in male, but not female, neurons in a sex hormone-independent, but mGluR5-dependent manner.

**Figure 4:**
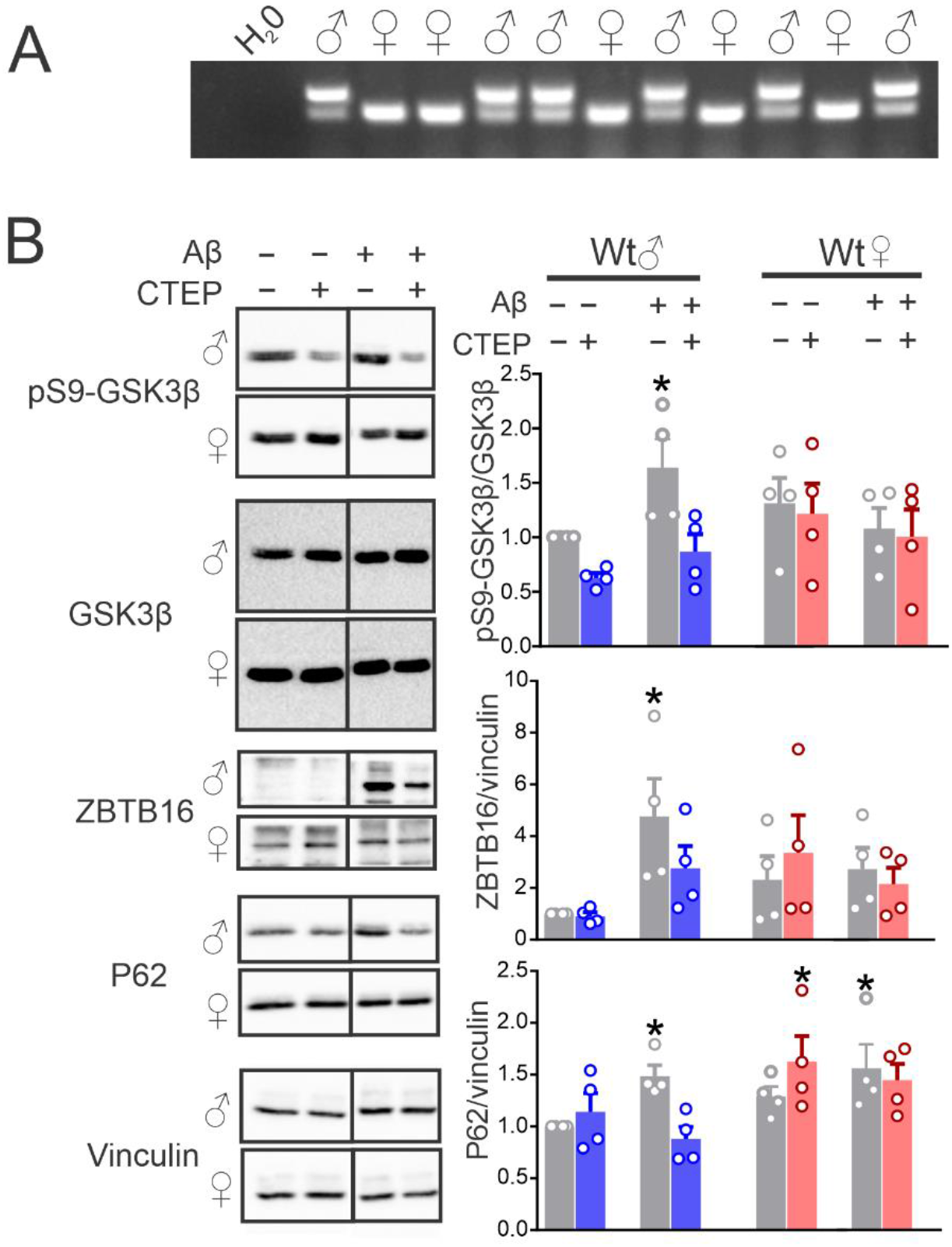
Sex-specific Aβ oligomer-mediated activation of mGluR5 signaling in neuronal cultures. **(A)** Representative gel for the Rbm31 gene PCR amplicon in male and female E18 wild-type (Wt) mouse embryos. **(B)** Representative immunoblots and quantification of pS9-GSK3β, ZBTB16 and p62 normalized to loading controls or total protein and expressed as a fraction of the untreated male cultures in primary cultured cortical neurons from male and female Wt E18 embryos stimulated with Aβ oligomers (100 nM) in the absence (DMSO) or presence of CTEP (10 μM). Data represents mean ± SEM (n=4 for each group). *P<0.05 versus untreated male cultures assessed by two-way ANOVA and Fisher’s LSD comparisons.

### mGluR5 inhibition reduced AD-related pathology in male, but not female, APP mice

We determined whether the differences in mGluR5-regulated GSK3β/ZBTB16 signaling between male and female cultured neurons after exposure to Aβ oligomers were translatable in vivo. To test this, 6 month old male and female wild-type and APP mice were treated with vehicle or CTEP (2 mg/Kg) for 12 weeks and brains were harvested at the end of treatment for biochemical and immunohistochemical assessments. We chose the APP model as both male and female mice exhibited a robust Aβ accumulation and cognitive impairment by 9 months *(28, 34)* and this treatment regime was effective in improving memory deficits and reversing Aβ pathology in 12-month-old male APP mice *(22)*. Here, we found that cell signaling mediators of the GSK3β/ZBTB16/ATG14 autophagic pathway were inhibited in male APP mice. Specifically, we detected an increased pS9-GSK3β level, increased ZBTB16 expression, reduced ATG14 expression and the accumulation of p62 protein (Fig. 5A-D) *(20, 32, 35)*. CTEP treatment of male APP mice attenuated pS9-GSK3β phosphorylation, reduced ZBTB16 protein expression, increased ATG14 protein expression resulting in a loss of p62 protein (Fig. 5A-D). When the same experiments were performed in female APP mice no alterations in GSK3β/ZBTB16/ATG14 signaling were observed in vehicle-treated female APP mice and as a consequence CTEP treatment had no discernable effect in on GSK3β/ZBTB16/ATG14 pathway in either wild-type or APP female mice (Fig. 5A-D).

**Figure 5:**
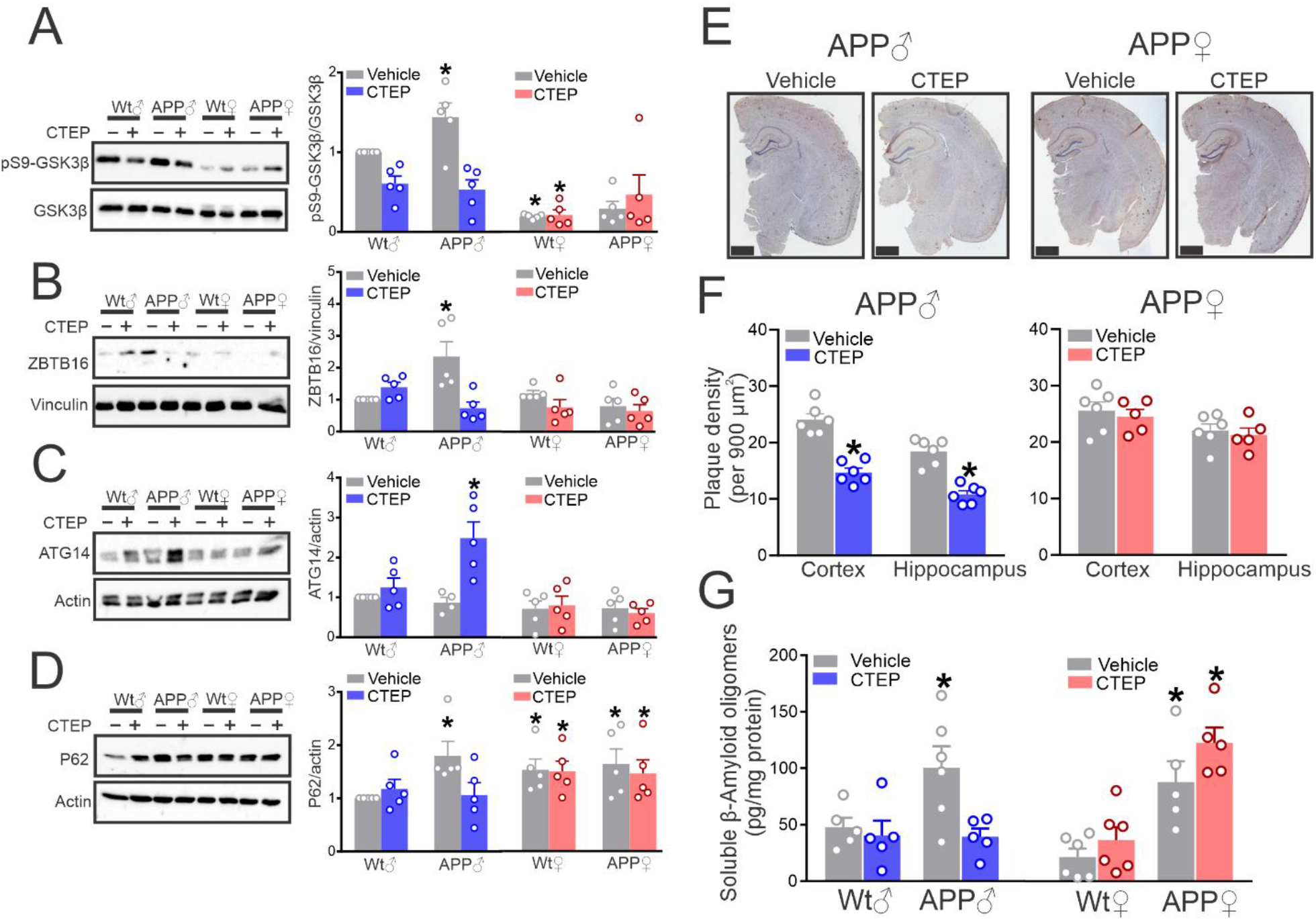
CTEP activates autophagy and reduces Aβ pathology in male, but not female, APP mice. Representative immunoblots and quantification of **(A)** pS9-GSK3β, **(B)** ZBTB16, **(C)** ATG14 and **(D)** p62 with the corresponding loading controls or total protein, **(E)** representative images of Aβ staining and **(F)** plaque density in cortical and hippocampal brain slices quantified of 5 different 900 μm^2^ regions from 6 brain slices per mouse (scale bar = 1 mm), and **(G)** Aβ oligomer concentrations (pg/mg) in 6 month old male and female wild-type (Wt) and APP mice treated with vehicle or CTEP (2 mg/kg) for 12 weeks. Data represents mean ± SEM (n=5 for each group) and quantification in **(A-D)** is expressed as a fraction of the vehicle-treated male Wt. *P<0.05 vs vehicle-treated male Wt values assessed by two-way ANOVA and Fisher’s LSD comparisons **(B-D and G)** or Kruskal-Wallis test **(A)**. *P<0.05 vs vehicle-treated same sex APP assessed by Student’s t-test **(F)**.

We then tested whether the sex-specific effect of CTEP on GSK3β/ZBTB16/ATG14 signaling in APP mice was correlated with a similar change in Aβ pathology. We found that the deposition of Aβ plaque density was significantly reduced in male APP mice following the CTEP treatment regime, whereas Aβ plaque density in either the hippocampus or cortex of female APP animals were not changed in response to CTEP treatment (Fig. 5E and 5F). Soluble Aβ levels detected in brain lysates from both vehicle-treated male and female APP mice were comparably increased versus sex-matched wild-type mice (Fig. 5G). CTEP treatment of male APP mice resulted in significantly reduced soluble Aβ levels, whereas it did not change soluble Aβ levels in female APP mice (Fig. 5G). Overall, these findings show that, while Aβ oligomers contribute to AD-like neuropathology in both sexes of APP mice, the Aβ oligomer-mediated antagonism of a mGluR5-regulated GSK3β/ZBTB16 autophagy contributing to Aβ pathology was exclusive to male mice. This may explain why mGluR5 inhibition reduced Aβ deposition in male APP mice only.

### mGluR5 blockade improved cognitive deficits in male, but not female, APP mice

We finally tested whether sex-specific outcomes of CTEP treatment on Aβ pathology were reflected on the memory function of APP mice. Vehicle and CTEP-treated male and female wild-type and APP mice were tested for impairments in working and spatial memory in the novel object recognition test and the Morris water maze (MWM) and Morris water maze with reversal (RMWM). In the novel object recognition test, vehicle-treated wild-type male and female mice discriminated between novel and familiar objects, whereas vehicle-treated male and female APP mice failed to discriminate between objects (Fig. 6A and 6B). Following CTEP treatment, male APP mice regained the capacity to discriminate between objects, whereas female APP mice remained cognitively impaired (Fig. 6A and 6B). In MWM and RMWM, vehicle-treated male and female APP mice exhibited significantly longer escape latencies and less time spent in target quadrant than did similarly treated same sex wild-type mice (Fig.7A-D). CTEP treatment improved male APP mice performance in both the MWM and RMWM as measured by shorter escape latency and longer time spent in target quadrant with values indistinguishable from wild-type mice (Fig. 7A-D). In contrast, when compared to vehicle-treated female APP mice, CTEP treatment did not improve either escape latency or time spent in target quadrant of female APP mice in either the MWM or RMWM (Fig. 7A-D). Unlike what we observed for wild-type male mice, CTEP treatment impaired female wild-type mouse performance in the RMWM compared to vehicle-treated wild-type mice, with the mice showing impaired escape latencies and reduced time spent in the target quadrant (Fig. 7C and 7D). Taken together, it is evident that male and female APP mice presented with comparable cognitive impairments and similar Aβ burdens, but the chronic inhibition of mGluR5 mitigated AD-like neuropathology in male, but not female, mice. These findings provide in vivo evidence for the sex-specific contribution of mGluR5 to AD pathophysiology and corroborate our primary neuronal cultures findings.

**Figure 6:**
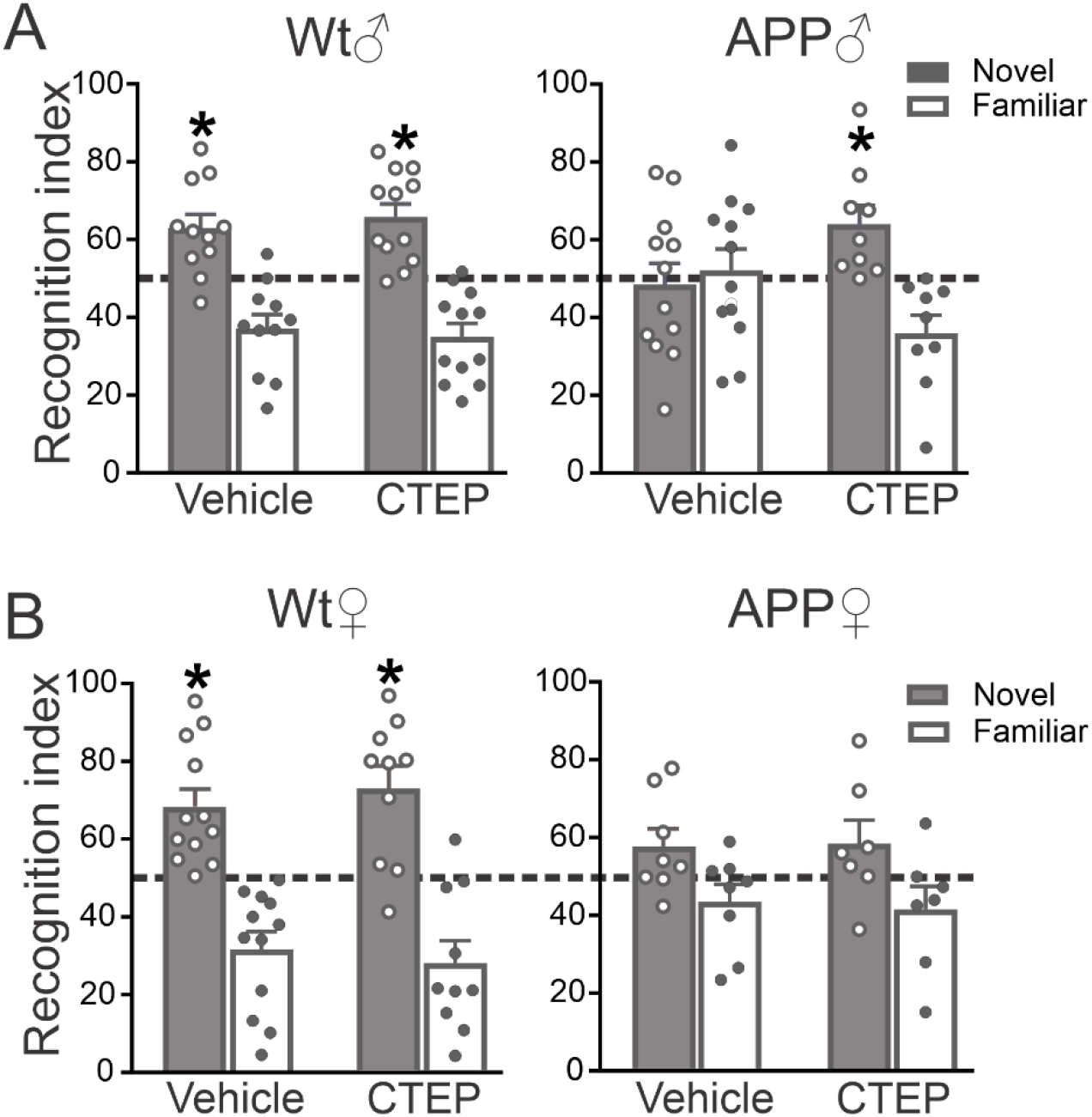
CTEP improves recognition scores in male, but not female, APP mice. Recognition index, for exploring one novel object versus familiar object in the second day of novel object recognition test following 12 week treatment with vehicle or CTEP (2mg/kg) of 6 month old **(A)** male **(B)** female wild-type (Wt) and APP mice. Data represents mean ± SEM (n=8-10 for each group). * P<0.05 versus familiar object assessed by two-way ANOVA and Fisher’s LSD comparisons. Mice were excluded from analysis due to spontaneous death

**Figure 7:**
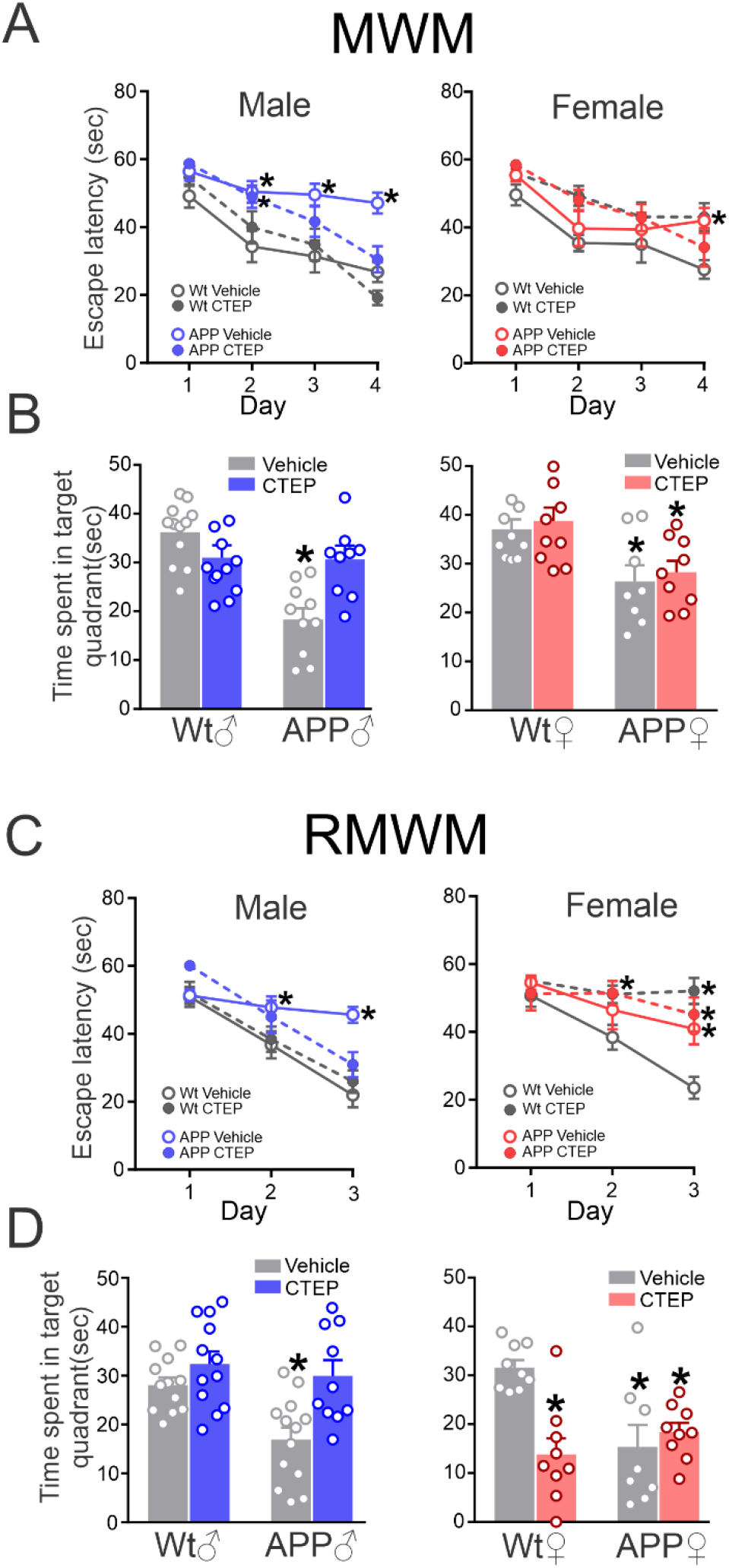
CTEP treatment improves performance of male, but not female, APP mice in MWM and RMWM. Escape latency and time spent in the target quadrant for MWM **(A and B)** and RMWM **(C and D)** in 6 month old male and female wild-type (Wt) and APP mice following 12 week treatment with either vehicle or CTEP (2 mg/kg). Data represents mean ± SEM (n=8-10 for each group). *P<0.05 vs vehicle-treated Wt assessed by two-way ANOVA and Fisher’s LSD comparisons. Mice excluded from analysis were due to spontaneous death.

## DISCUSSION

One major challenge in AD therapy is the identification of a pharmaceutical target that both manages disease symptomology and modifies disease progression. Genetic and pharmacological silencing of mGluR5 has identified a contributory role for mGluR5 in Aβ-related cognitive decline and pathology in male AD mice *(21, 22)*. However, gender-specific responses to therapies have been reported in many neurological diseases including AD which have the potential to further complicate the process of drug discovery *(7, 8, 10)* and therefore, delineation of sex-specific differences in AD pathophysiology became essential. We show here that, unlike what is observed for mGluR5 expressed in male tissue, mGluR5 is incapable of scaffolding either Aβ oligomers or PrP^C^ in female mouse brain tissue to elicit pathological mGluR5 signaling. This failure to form a mGluR5-scaffolded complex is conserved in both mouse and human female cortical tissue and represents an unexpected, but novel sex-specific regulation of mGluR5 pharmacology and signaling that appears to be evolutionarily conserved. The sex-dependent contribution of mGluR5 to AD pathology was then validated in vivo since treatment of APP mice with a mGluR5 NAM improved cognitive deficits and reduced Aβ-related pathology in males, but was ineffective in females despite a comparable disease phenotype. This study has major implications as it shows how GPCRs, the most common class of receptors targeted by prescription drugs *(36)*, can exhibits district gender-specific pharmacological profiles that must be considered during the process of drug development. Moreover, it highlights the need for sex-specific stratification of neurodegenerative drug trial results that will impact the guidance of future design of sex-tailored treatment strategies.

The ability of PrP^C^ and Aβ oligomers to form a scaffolding complex with male mGluR5 can promote receptor clustering at the synapses *(17, 19)* and prevent the constitutive receptor endocytosis *(37)* is crucial in activating the pathological signaling of mGluR5 *(5, 38)*. One of the consequences of pathological mGluR5 activation by Aβ is the inhibition of autophagy flux *(20, 24)* that can then trigger a feedforward mechanism leading to further accumulation of Aβ oligomers and exacerbation of glutamatergic excitotoxicity. mGluR5 NAM via its binding to the allosteric site of mGluR5, may disrupt the interaction between Aβ oligomers and the receptor and consequently interrupt receptor-activated neurodegeneration. In fact, we have previously reported that mGluR5 NAM treatment normalizes the increased cell surface expression and signaling of mGluR5 in male APP mice *(20)*. The lack of PrP^C^/Aβ oligomer binding to female mGluR5 means that this feedforward mechanism cannot be initiated by mGluR5 and therefore nullify the contribution of mGluR5 to AD pathophysiology in female AD mice and explains the lack of mGluR5 NAM efficacy in female mice.

Aβ oligomers represent a normal proteolytic byproduct of amyloid precursor protein metabolism and are normally found at low levels in the brain. Alterations in the production and/or clearance of Aβ oligomers can shift the homeostasis towards increased cellular Aβ oligomer levels and accelerate neurodegeneration *(4, 39)*. Therefore, enormous effort has been directed toward identifying novel pathways that can regulate Aβ oligomer accumulation. Within this context, autophagy represents an essential protein degradation pathway and defects in autophagy were reported in the early stages of AD in both animal models and patients *(40–44)*. We find that mGluR5 triggers pathological inhibition of autophagy via a ZBTB16-Cullin3-Roc1 E3-ubiquitin ligase pathway *(35)* in male, but not female, APP mice. The antagonism of mGluR5 was associated with improved cognitive function and reduced Aβ pathology in male APP mice, but unfortunately this improvement was not observed in female mice.

We and others have previously reported that both mGluR5 negative and silent allosteric modulators improve cognitive function in male APP mice, but only NAMs reduce Aβ oligomer-related pathology *(22, 25)*. This combined data suggests the possibility of repurposing mGluR5 selective modulators for treatment of male AD patients. This is of particular interest since the mGluR5 NAM basimglurant has been shown to be well-tolerated by patients in clinical studies for Fragile X mental retardation and major depressive disorder *(45, 46)*. However, a critical drawback for repurposing mGluR5 selective drugs to treat AD in the general population is that mGluR5 fails to interact with Aβ oligomers in female tissue and therefore, mGluR5 contribution to cognitive decline and AD pathology in females is not significant and must be mediated by alternative mechanism(s). Nevertheless, important opportunities still exist for the treatment of AD in men by mGluR5-selective drugs.

Our a priori expectation predicted that mGluR5 antagonism would be equally effective in both males and females, as there have been no reported differences in either gene editing or genomic regulation of male versus female mGluR5 expression, subcellular localization and/or function. Thus, one of the major challenges of AD research remains understanding the underlying pharmacological and physiological differences in the manifestation of AD pathology in males versus females, in both animal AD mice and human patients. The sex-dependent regulation of the formation of the mGluR5/Aβ oligomer/PrP^C^ complex is clearly one of the underlying mechanisms for why mGluR5 antagonism is not effective in the treatment of cognitive impairment and Aβ pathology in female mice. However, this leaves open the question as to what is the underlying biological reason for why mGluR5 exhibits differential pharmacological properties in neurons of both sexes. Metabotropic glutamate receptors are allosterically regulated receptors and thus their pharmacological function may be regulated by differential interactions with either intra- or extracellular regulatory proteins in male and female animals *(16, 47)*. Moreover, we cannot rule out sex-specific difference in either mGluR5 variant splicing or interactomes that may influence its binding to other extracellular scaffolds such as PrP^C^ and Aβ.

The pharmacology underlying disease etiology and progression regulated by cellular processes that are independent of mGluR5 in females remains to be determined. The important underlying message from the current study is that mGluR5 is not a contributor to disease progression and pathology in female mice, thereby severely limiting the use of mGluR5 antagonists in women. However, it is important to acknowledge that mGluR5 is not the only cell surface target for Aβ oligomers and a multicity of different targets can contribute to sex-dependent and -independent changes in synaptic signaling, synaptic pruning, autophagy and neuronal cell death in female AD *(48)*. Since we detect impairment in autophagy flux in female AD mice, it will be also important to explore specific alterations in the other autophagic signaling mechanism(s) in female AD brain and identify novel approaches to effectively target them. More so, given the role of mGluR5 in synaptic function *(5, 49)* and the poor cognitive performance of mGluR5 NAM-treated control female mice, it will be essential to exploit other aspects of altered mGluR5 synaptic signaling in female brain.

In summary, we report that sex-specific differences in mGluR5 pharmacology observed in mouse cortical tissue are mirrored in human cortex and show that mGluR5 antagonism reduces cognitive impairment and Aβ pathology in male, but not female, AD mice. Our data clearly demonstrates that mGluR5 does not represent an effective pharmacological target for the treatment of AD in women, but that it still remains a potentially effective target for the treatment of the early stages of AD in men. This study highlights the need for the sexual stratification of neurodegenerative drug trial result to aid in the development of sex-specific therapeutic strategies.

## MATERIALS AND METHODS

### Reagents

CTEP (1972) was purchased from Axon Medchem. β-Amyloid [1 42] PTD Human protein (03111), Amyloid beta (Aggregated) Human ELISA Kit (KHB3491), Sulfo-NHS-SS-Biotin (21331), NeutrAvidin Resins (29200), goat anti-Rabbit (G-21234) and anti-mouse (G21040) IgG (H+L) Cross-Adsorbed HRP Secondary Antibody, rabbit anti-β-Amyloid (71-5800) and rabbit anti-β-Actin (PA1-183) were from Thermo Fisher Scientific. Rabbit anti-pS9-GSK3β (9323) and mouse anti-GSK3β (9832) antibodies were from Cell Signaling Technology. Rabbit anti-ATG14L (PD026) was from MBL International. Immunoprecipitation kit (206996), mouse anti-P62 (56416), rabbit anti-vinculin (129002) and rabbit anti-ZBTB16 (39354) were from Abcam. Rabbit anti-mGluR5 (AB5675) was from Millipore. Mouse Anti-Prion Protein (PrP^C^) Clone SAF-32 (189720-1) was from Cayman Chemical. [^3^H] MPEP (VT237) was from Vitrax, MPEP (1212) was from Tocris, Effectene Transfection reagent (301425) was from Qiagen and VECTASTAIN Elite ABC HRP Kit (Rabbit IgG, PK-6101) was from Vector Laboratories. Reagents used for western blotting were purchased from Bio-Rad and all other biochemical reagents were from Sigma-Aldrich.

### Animals

Mice were obtained from The Jackson laboratories and bred to establish littermate controlled female and male wild-type (C57BL/6J, stock# 000664), APPswe/PS1ΔE9 (B6C3-Tg (APPswe/PSEN1ΔE9)85Dbo/J, stock# 34829) and mGluR5^−/−^ (B6;129-Grm5tm1Rod/J, stock# 003121) that were group-housed in cages of 2 or more animals, received food and water ad libitum and maintained on a 12-hour light/12hour dark cycle at 24°C. Groups of 24 male and female Wt and APPswe/PS1ΔE9 mice were aged to 6 months of age and 12 mice from each group were randomized and blindly-treated every 48h based on weekly weights with either vehicle (DMSO in chocolate pudding) or CTEP (2 mg/kg, dissolved in 10% DMSO then mixed with chocolate pudding, final DMSO concentration was 0.1%) for 12 weeks *(22, 32)*. Cognitive function of all animals was assessed prior to and following 12 weeks of drug treatment. At the end of the 12-week treatment, mice were sacrificed by exsanguination and brains were collected and randomized for biochemical determinations and immunostaining. Mice used for radioligand binding and coimmunoprecipitation experiments were 3-month-old Wt. All animal experimental protocols were approved by the University of Ottawa Institutional Animal Care Committee and were in accordance with the Canadian Council of Animal Care guidelines.

### Human specimens

All tissues were obtained collected after consent was provided from patients or the next kin in accordance with institutional review board-approved guidelines. Frozen samples of cortices from subjects under 50 years of age were acquired through the University of Alabama and the Autism Tissue Program. Additional, frozen specimens from adults above age 60 years were obtained from the Neuropathology Service at Brigham and Women’s Hospital and the Department of Pathology and Laboratory Medicine at The Ottawa Hospital. Patients’ gender, age and diagnosis were as follow: male, 34-year-old, vasculitis/encephalitis; male, 70-year-old, dementia with lewy bodies; male, 65-year-old, dementia with lewy bodies; male, 75-year-old, healthy; female, 44-year-old, carcinoma; female, 55-year-old, healthy; female, 65-year-old, healthy; female, 73-year-old, healthy.

### Cell lines

CF10 cells were maintained in Opti-MEM supplemented with FBS (10% v/v), respectively at 37 °C in 5% CO_2_ humidified incubator. CF10 cells were transfected with FLAG-mGluR5 ± PrP^C^ using Effectene transfection reagents following the manufacture’s protocol.

### Primary neuronal culture

Cultures were prepared from the cortical region of E18 male and female wild-type embryo brains. Briefly, cortical tissue of each mouse embryo was trypsin-digested followed by cell dissociation using a fire-polished Pasteur pipette. Cells were plated on Poly-L-Ornithine-coated dishes in neurobasal medium supplemented with N2 and B27 supplements, 2.0 mM GlutaMAX, 50 μg/ml penicillin, and 50 μg/ml streptomycin. Cells were maintained for 12 to 15 days at 37 °C in 5% CO_2_ humidified incubator before experimentation.

## Experimental Methods

### Novel object recognition

Mice were habituated in the testing room for 30 mins and testing was blindly performed during the animal’s light cycle. Mice were placed in the empty box measuring 45 × 45 × 45 cm for 5 min and 5 min later, 2 identical objects were placed in the box 5 cm from the edge and 5 cm apart. Mice were returned to the box for 5 min, and allowed to explore, as described previously *(32)*. Time spent exploring each object was recorded using a camera fed to a computer in a separate room and analyzed using Noldus Ethovision 10 software. Mice were considered to be exploring an object if their snout was within 1 cm of the object. Each experiment was repeated 24 hours after first exposure with one object replaced with a novel object. Data was interpreted using recognition index which was as follows: time spent exploring the familiar object or the novel object over the total time spent exploring both objects multiplied by 100, and was used to measure recognition memory (TA or TB/ (TA + TB))*100, where T represents time, A represents familiar object and B, novel object.

### Morris water maze (MWM) and reversal Morris water maze (RMWM)

Animals were habituated in the testing room for 30 min and testing was blindly performed during the animal’s light cycle. The Morris water maze test was performed in a white opaque plastic pool (120 cm in diameter), filled with water and maintained at 25 °C to prevent hypothermia, as described previously *(22)*. A clear escape platform (10 cm diameter) was placed 25 cm from the perimeter, hidden one cm beneath the surface of the water. Visual cues were placed on the walls in the room of the maze as spatial references. Mice were trained for 4 days (four trials per day and 15 mins between trails) to find the submerged platform at a fixed position from a random start point of the 4 equally spaced points around the pool. Each trial lasted either 60 seconds or until the mouse found the platform and mice remained on the platform for 15 seconds before being removed to their home cage. If the mice failed to find the platform within 60 seconds, they were guided to the platform by the experimenter. Escape latency was measured using Ethovision 10 automated video tracking software from Noldus. On day 5, the probe trial (a single trial of 60 seconds) was performed by removing the platform and allowing the mice to swim freely in the pool and recording the time spent in the target quadrant. RMWM task was initiated 24 hours after completion of MWM using the same paradigm as MWM, with 4 days acquisition and probe trial. In the RMWM task, the platform was relocated to a new position.

### Sandwich ELISA for Aβ oligomer levels

ELISA was performed as described previously *(22, 24)*. Briefly, brains were dissected and one hemisphere was used to analyze oligomeric Aβ levels. Brain homogenates were divided and centrifuged at 4°C at 100,000 ×g for 1 hour. The supernatant was then diluted 1:10 with kit-provided buffer before carrying out the ELISA, which was performed in triplicate and measured as detailed in the manufacturer’s protocol. Protein concentrations were quantified using the Bradford protein assay (Bio-Rad). The final Aβ concentrations were determined following normalization to total protein levels.

### β-Amyloid immunohistochemistry

Immunostaining was performed as described previously *(22, 24)*. Briefly, brains were coronally sectioned through the cortex and hippocampus and staining was performed on 40 μm free-floating sections using a peroxidase-based immunostaining protocol (VECTASTAIN Elite ABC HRP Kit). Sections were incubated overnight in primary antibody for Aβ (1:200) at 4 °C, washed, incubated in biotinylated antibody (biotinylated horse anti-rabbit, 1:400) for 90 mins at 4 °C, then incubated in an avidin biotin enzyme reagent for 90 min at 4°C and visualized using a chromogen. Sections were mounted on slides and visualized with a Zeiss AxioObserver epifluorescent microscope with a Zeiss 20× lens, using representative 900 μm^2^ areas of cortex and hippocampus. Experimenters were blinded to drugging and analysis. Six to eight sections per mouse were analyzed and for each section 5 ROIs were analyzed in the cortex and 2 ROIs in the hippocampus using the cell counter tool in Image J *(50)*. This number of ROIs prevents the selection of only densely stained regions.

### Immunoblotting

A brain hemisphere or neuronal culture dish was lysed in ice-cold lysis buffer (50 mM Tris, pH 8, 150 mM NaCl, and 1% Triton X-100) containing protease inhibitors cocktail (100 μM AEBSF, 2 μM leupeptin, 80 nM aprotinin, 5 μM Bestatin, 1.5 μM E-64 and 1 μM pepstatin A) and phosphatase inhibitors (10 mM NaF and 500 μM Na_3_VO_4_) and centrifuged twice for 10 min each at 20,000 xg and 4 °C. The supernatant was collected and total protein levels were quantified using Bradford Protein Assay. Homogenates were diluted in a mix of lysis buffer and β-mercaptoethanol containing 3x loading buffer and boiled for 10 min at 95 °C. Aliquots containing 30-40 μg total proteins were resolved by electrophoresis on a 7.5% or 10% SDS-polyacrylamide gel (SDS-PAGE) and transferred onto nitrocellulose membranes. Blots were blocked in Tris-buffered saline, pH 7.6 containing 0.05% of Tween 20 (TBST) and 5% non-fat dry milk for 2 h at room temperature and then incubated overnight at 4 °C with primary antibodies diluted 1:1000 in TBST containing 2% non-fat dry milk. Immunodetection was performed by incubating with secondary antibodies (anti-rabbit/mouse) diluted 1:5000 in TBST containing 1% of non-fat dry milk for 1 h. Membranes were washed in TBST and then bands were detected and quantified using SuperSignal™ West Pico PLUS Chemiluminescent Substrate using Bio-Rad chemiluminescence system as previously described *(32, 51)*.

### Embryo sex determination for primary culture

DNA was extracted from embryonic tissue by incubation overnight at 55 °C in DNA lysis buffer (50 mM Tris-HCl, pH 8, 10 mM EDTA, 20 mM NaCl and 0.031% SDS) supplemented with Proteinase K. After incubation, lysates were centrifuged at 20,000 xg for 10 min at 25 °C and the supernatant was collected. DNA was precipitated by Isopropanol and was collected by centrifugation at 10,000 xg for 10 min at 4 °C. Tris-EDTA buffer (10 mM Tris-HCl, pH 8 and 1 mM EDTA, pH 8) was added to DNA, heated at 55 °C for 10 min and then used for the PCR reaction. Primers were designed flanking an 84 bp deletion of the X-linked Rbm31x gene relative to its Y-linked gametolog Rbm31y (Forward; CACCTTAAGAACAAGCCAATACA and Reverse: GGCTTGTCCTGAAAACATTTGG) *(33)*. Following the PCR reaction, products were separated on an agarose gel containing RedSafe DNA stain (FroggaBio) and visualized on Bio-Rad fluorescence system.

### Primary neuronal culture experiment

Following 12-15 days incubation, cultures were starved in HBSS for 1h. Cells were then treated with 10 μM CTEP or DMSO (vehicle for CTEP) for 30 min followed by 100 nM Aβ for 1 h at 37 °C. Human Aβ oligomers were prepared as per the manufacturer’s recommendations to form the neurotoxic aggregates *(52)*. Following this treatment, neuronal cultures were lysed with ice-cold RIPA buffer supplemented with protease and phosphatase inhibitors then immunoblotting was performed.

### Radioligand binding

Radioligand binding was performed as previously described *(53)*. Briefly, cells washed with PBS and ice-cold lysis buffer (10 mM Tris-HCl, pH 7.4 and 5 mM EDTA, pH 8.0) was used to detach the cells. Cortex from mice and human were homogenized in ice-cold lysis buffer. To prepare the crude membrane preparation, cell or brain lysates were centrifuged twice at 40,000 xg for 20 min at 4 °C and supernatant was discarded after each centrifugation and the pellets were resuspended in lysis buffer. The crude membrane preparations were then suspended in resuspension buffer (62.5 mM Tris-HCl, pH 7.4 and 1.25 mM EDTA, pH 8.0) and kept on ice. The binding reactions were performed in duplicate by incubating crude membranes with 3 nM [^3^H]-MPEP and increasing concentrations of either MPEP or Aβ oligomers in binding buffer (62.5 mM Tris-HCl, pH 7.4 and 1.25 mM EDTA, pH 8.0, 200 mM NaCl, 6.7 mM MgCl_2_, 2.5 mM CaCl_2_ and 8.33 mM KCl) at room temperature for 90 min. The nonspecific binding was determined using 10 μM MPEP. The binding reactions were terminated by rapid filtration through Whatman GF/C glass fiber filter sheets using semi-automated harvesting system (Brandel). The tritium-bound radioactivity was then counted using liquid scintillation counter (Beckman). The data were analyzed using GraphPad Prism, [^3^H]-MPEP binding as % of maximum specific binding was calculated and the equilibrium dissociation constant (K_i_, M) of MPEP and Aβ were determined.

### Co-immunoprecipitation

Hippocampus from male and female wild-type and mGluR5^−/−^ mice were dissected and lysed in non-denaturing lysis buffer containing protease inhibitors provided with co-immunoprecipitation kit. Lysates were rotated for 1 hour at 4 °C and centrifuged at 10,000 g for 10 min at 4 °C to pellet insoluble material. Precleared supernatant (500μg) was incubated with 1 μg of anti-PrP^C^ antibody over night at 4 °C. Freshly washed protein A/G-sepharose beads were added to lysate/antibody mixture and samples were rotated for 2 hours at 4 °C. Beads were washed with wash buffer provided in the kit and then boiled with 3x loading buffer containing β- mercaptoethanol for 10 min at 90 °C. Samples were separated by SDS-PAGE and immunoblotted to identify co-immunoprecipitated mGluR5. An additional immunoblot was performed to examine mGluR5 and PrP^C^ protein expression in lysates prepared before incubation with antibody.

### Statistical analysis

Means ± SEM are shown for each of independent experiments are shown in the various figure legends. GraphPad Prism 8 was used to analyze data for normality and statistical significance. Data normality was tested using Anderson-Darling and D’Agostino-Pearson omnibus tests and statistical significance was determined by two-way ANOVA, Student’s t test or Kruskal-Wallis test as appropriate for the significant main interactions. Statistical details of individual experiments are indicated in figure legends

**Figure 8:**
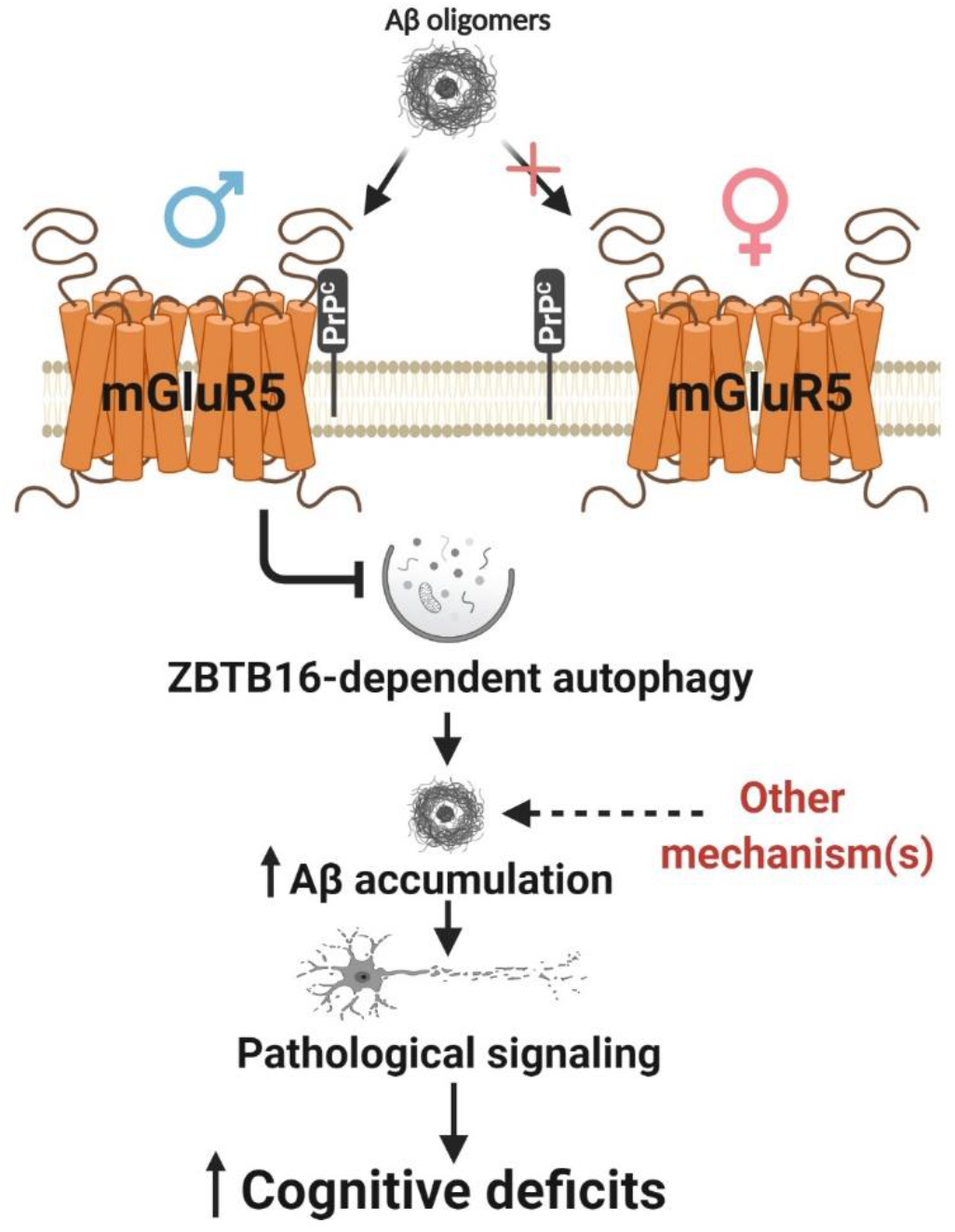
Schematic representation depicting Aβ oligomer induced sex-selective pathophysiological mGluR5 signaling. PrP^C^ forms a scaffolding complex with male mGluR5 and Aβ oligomers and elicit the inactivation of ZBTB16-mediated autophagy. The reduction in autophagic clearance of Aβ oligomers triggers pathological signaling and results in cognitive deficits. In female brain, lack on interaction between PrP^C^ and mGluR5 prevents the formation of mGluR5/Aβ oligomer/PrP^C^ complex and therefore pathological signaling and cognitive deficits are likely mediated by mechanism(s) other than mGluR5.

## ACKNOWLEDGMENTS

S.S.G.F is a Tier I Canada Research Chair in Brain and Mind. K.S.A is a Lecturer at the Department of Pharmacology and Toxicology, Faculty of Pharmacy, Alexandria University. Thanks to Shaunessy Hutchinson for breeding the mouse colony, the Behavior and Physiology core at the University of Ottawa, Bassam Albraidy for technical assistance, Dr. Suzette Priola (NIH) for providing CF10 cells, Dr. Andrew West (Duke University), Dr. Jennifer Chan (University Calgary) and Dr. John Woulfe (The Ottawa Hospital) for their assistance in procuring human tissues.

## FUNDING

This study was supported by Canadian Institutes for Health Research (CIHR) grants (PJT-148656 and PJT-165967) and Funding from Krembil Foundation to S.S.G.F, and a Clinician Postdoctoral Fellowship from the Alberta Innovates Health Solutions (AIHS) and CIHR to K.S.A.

## AUTHOR CONTRIBUTIONS

Conceptualization, K.S.A., A.H. and S.S.G.F; Methodology, Formal analysis and investigation, K.S.A., A.A., J.M.D. and A.H.; Resources, F.M.R., M.G.S., M.T., S.S.G.F.; Writing-Original draft, K.S.A.; Writing-Review & Editing, K.S.A., S.S.G.F.; Supervision and Funding acquisition, S.S.G.F.

## COMPETING INTERESTS

Authors declare no conflict of interest.

### DATA AND MATERIALS AVAILABILITY

The published article includes all datasets generated and analyzed during this study.

## Notes

### Competing Interest Statement

The authors have declared no competing interest.

### Summary of Updates

The abstract and title were revised to better reflect the findings of the study and results were expanded to better explain the data.

